# Sleep-specific changes in physiological brain pulsations

**DOI:** 10.1101/2020.09.03.280479

**Authors:** H Helakari, V Korhonen, SC Holst, J Piispala, M Kallio, T Väyrynen, N Huotari, L Raitamaa, J Tuunanen, J Kananen, M Järvelä, V Raatikainen, V Borchardt, H Kinnunen, M Nedergaard, V Kiviniemi

**Author notes:** Corresponding Author Vesa Kiviniemi, Prof, MD, Department of Diagnostic Radiology, Medical Research Center (MRC), Oulu University Hospital, Kajaanintie 50, 90220, Oulu, Heta Helakari, MSc, Department of Diagnostic Radiology, Medical Research Center (MRC), Oulu University Hospital, Kajaanintie 50, 90220, Oulu.

## Abstract

Sleep is known to increase the convection of interstitial brain metabolites along with cerebrospinal fluid (CSF). We used ultrafast magnetic resonance encephalography (MREG_BOLD_) to quantify the effect of sleep on physiological (vasomotor, respiratory and cardiac) brain pulsations driving the CSF convection in humans. Transition to electroencephalography verified sleep occurred in conjunction with power increase and reduced spectral entropy (SE) of physiological brain pulsations. During sleep, the greatest increase in spectral power was in very-low frequency (VLF < 0.1 Hz) waves, followed by respiratory and cardiac brain pulsations. SE reduction coincided with decreased vigilance in awake state and could robustly (ROC 0.88, p < 0.001) differentiate between sleep vs. awake states, indicating the sensitivity of SE of the MREG_BOLD_ signal as a marker for sleep level. In conclusion, the three physiological brain pulsation contribute to the sleep-associated increase in glymphatic CSF convective flow in an inverse frequency order.

**Highlights:**

- Brain tissue contains almost no connective tissue, this enabling pressure waves to initiate long-distance brain pulsations
- Brain pulsations are induced by vasomotion, respiration, and the cardiac cycle
- Sleep strikingly increases spectral power and decreases spectral entropy of brain pulsations, especially for the very low frequency vasomotor waves
- Spectral entropy of brain pulsations detected by MREG is a sensitive measure of vigilance, resembling the corresponding entropy changes detected by scalp EEG

## Introduction

A rarely considered feature of the mammalian brain is that its neural tissue consists of almost 80% water, with a near absence of connective tissue. Combined with the physical constraint of floating in cerebrospinal fluid (CSF) embedded within an incompressible cranial vault, physiological pressure changes can thus drive brain-wide pulsations and fast fluid movement within the neuropil. In the early 20^th^ century studies of pulsatile brain activity that lead to the invention of electroencephalography (EEG) for external detection of electrophysiological brain activity, Hans Berger also described three forms of brain pressure pulsations: “eine pulsatorische, eine respiratorische und vasomotorische Bewenung” (Berger, 1901) (pulsatory, respiratory and vasomotor movements). Nearly a century later, the same triad of cardiovascular, respiratory and slow, spontaneous vasomotor waves driven by arteries, can be non-invasively imaged throughout brain by magnetic resonance encephalography (MREG), a ultrafast variant of functional magnetic resonance imaging (fMRI) (Kiviniemi et al., 2016). MREG was invented specifically to maximize temporal accuracy while still maintaining adequate image quality for high frequency mapping of the fMRI blood oxygen level dependent (BOLD) signal.

Prior reports in rodents showed that natural sleep and certain anesthetic regiments can induce a sharp increase in the influx of CSF to brain tissue, suggesting a transition to a state more permissive to fluid dispersion (Xie et al., 2013). Human EEG studies link increased slow wave activity during sleep with decreases in cerebral blood volume that are consistently followed by greater influx of CSF into the fourth ventricle (Fultz et al., 2019). Insofar as EEG is a marker of net neural activity, its association with CSF movement is most conspicuous during sleep, when brain activity is relatively quiescent. Given this background, we tested the hypothesis that MREG imaging could reveal consistent changes in fluid pulsation within the neuropil during the transition of healthy human subjects from wakefulness to natural sleep.

In support of our hypothesis, we observed striking increases in the brain pulsation induced by each of the three types of pressure waves as the subjects entered an EEG-verified NREM sleep state. Power spectrum analysis revealed that the rank order of the enhanced brain pulsations during NREM sleep were slow vasomotion > respiration > cardiac cycle Similar to analogous EEG results acquired during sleep (Burioka et al., 2005; Liang, Kuo, Hu, Pan, & Wang, 2012; Mahon, Greene, Lynch, McNamara, & Shorten, 2008; Rodríguez-Sotelo et al., 2014; Vakkuri et al., 2005), the entropy of the MREG brain pulsation power spectrum was consistently and specifically decreased during NREM sleep, thus allowing higher sensitivity in the prediction of sleep state than are afforded by scoring of EEG recordings in 30 s epochs based on American Academy of Sleep Medicine (AASM) guidelines for clinical sleep recordings. Taken together, these observations confirm that the hydrostatic properties governing CSF movement in the brain are potently regulated across the sleep-wake cycle.

## Results

Due both to technical reasons and historical convention, the hemodynamic BOLD signal (< 0.5 Hz) and electrophysiological EEG (> 0.5 Hz) signals have generally been measured in non-overlapping spatiotemporal windows. At present, technical advances enable synchronous high density direct-current (DC)-EEG to be measured synchronously with ultrafast 10 Hz MREG_BOLD_ signal, which allows simultaneous whole brain assessment of hemodynamic and electrophysiological brain activity from temporally overlapping window extending from 0 to 5 Hz (Hiltunen et al., 2014; Keinänen et al., 2018; Richards, Boswell, Stevens, & Vendemia, 2015). Figure 1 depicts an example of sleep induced changes in the full 5 Hz band brain pulsations captured by simultaneous multimodal MREG and DC-EEG scanning over this wide range of low frequencies is illustrated in Figure 1.

**Figure 1.**
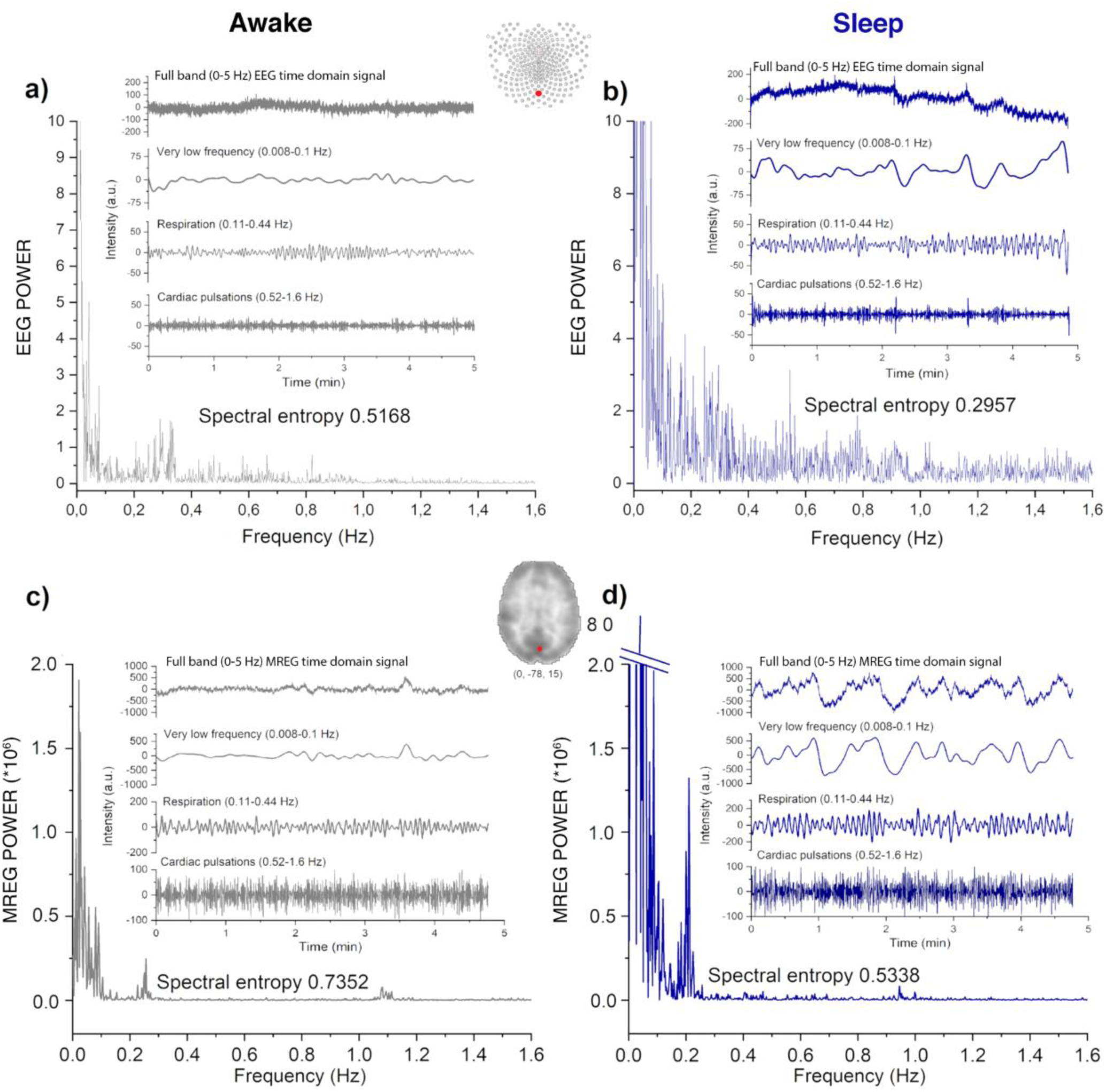
Examples of electrophysiological and dynamic brain pulsations measured in the same subject while awake and sleeping. The data was measured simultaneously by a-b) 256 lead high density DC-EEG (Oz) and c-d) ultrafast 10 Hz MREG_BOLD ._covering whole brain in 0 - 5 Hz. Time and frequency domain data both indicate increased power and amplitude of brain pulsations upon transition from EEG-verified fully awake state to fully NREM sleep. The dominant increase in VLF pulsation power reshapes the power spectrum distribution and alters the entropies of the EEG and MREG_BOLD_ signals.

### Spectral entropy of MREG signal predicts NREM sleep

Based on pilot evidence for the successful measurement synchronous multimodal MREG and DC-EEG signal (Fig. 1), we recruited healthy subjects for a sleep monitoring study lasting an entire week. Subjects were scanned after a normal night’s sleep and (three days later) after a monitored interval of wakefulness lasting 24 ± 1.5 h. The sleep deprivation was used to induce drops in awake vigilance for multimodal monitoring of associated changes in physiological brain pulsations (Fig. 2a). After sleep deprivation, the subjects exhibited significantly increased reaction times, indicative of reduced vigilance (CANTAB RTI test: 427 ± 25 > 403 ± 29 msec, p=0.002) compared to neuropsychology test results during an awake scan following a normal night’s sleep (Supplementary Fig.1).

**Figure 2.**
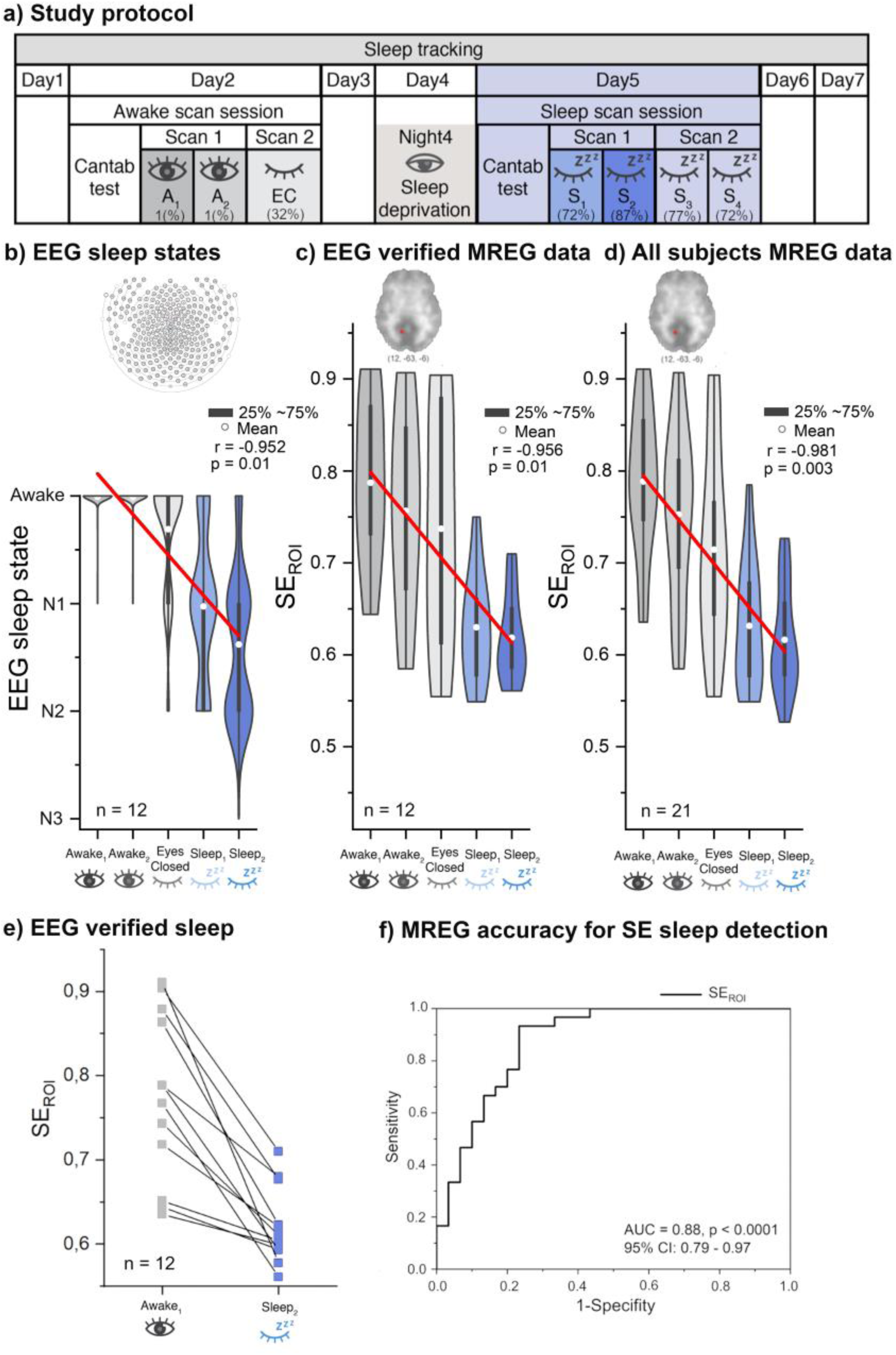
a) Study protocol shows overall study design through one week of sleep monitoring. Awake scanning session was arranged on a day two after a good night of sleep. Sleep scanning session was arranged on day five, after a night of sleep deprivation on the preceding night. b) EEG sleep state data show that the amount and depth of sleep both increase as a function of scan epoch. Visual cortex SE_ROI_ of MREG_BOLD_ data predicts sleep and wakefulness across subjects both in c) EEG-verified cases (n = 12) and d) all for all cases (n = 21)) as a function of scan epoch, resembling corresponding results for EEG-verified sleep states. Results indicate linear declines both in EEG sleep state and MREG_BOLD_ SE_ROI_ values as a function of scan epoch in the experiment. The amount of NREM sleep also increased in the successive scans, based on EEG-verified data. e) The SE_ROI_ showed a clear drop in all subjects in the transition from waking to EEG-verified sleep states S2. f) The ROC curve of SE_ROI_ data indicates high accuracy AUC = 0.88 (p < 0.0001) in the ability to separate sleep (n = 30 sleep epoch scans) from awake data (n = 30 epochs), an effect also observed in the whole brain SE_GLOBAL_ signal (supplementary Fig 2.).

Spectral entropy (SE) of brain EEG signals is known to reflect vigilance, sleep and depth of anesthesia (Burioka et al., 2005; Liang et al., 2012; Rodríguez-Sotelo et al., 2014). We compared the fitness of SE in MREG_BOLD_ data versus SE in simultaneously measured EEG recordings to predict vigilance. The EEG data were analyzed and scored as awake or sleep according to standard AASM EEG sleep scoring criteria. The SE evaluated over the whole brain (SE_GLOBAL_) revealed a negative linear relationship (r_pearson_=-0.35; p = 0.002, degrees of freedom, df 29, Supplementary Fig. 2a) with EEG-weighted sleep scoring as a function of scan epochs. We focused our analysis on the corresponding of MREG_BOLD_ SE in a region of interest (SE_ROI_) in the right visual cortex, which we selected due to the from known changes in posterior VLF pulsation during sleep (Chang et al., 2016; Liu et al., 2018). This comparison indicated a nearly two-fold stronger correlation for MREG_BOLD_ SE with the EEG-weighted sleep score (r_pearson_=-0.6, p < 0.001, df 29, Supplementary Fig. 2b). Furthermore, the AASM EEG sleep score data in EEG-verified scan epochs showed a similar linear relationship to the amount of sleep (r = -0.952, p=0.01) as did the individual MREG_BOLD_ SE_ROI_ (r = -0.956, p=0.01) and SE_ROI_ measured from all subjects (r = -0.981, p=0.003), thus confirming the fitness of SE_ROI_ as a measure of sleep (Figs. 2b-e). The accuracy by which the SE_ROI_ separated sleep from wakefulness episodes in MREG_BOLD_ data was further tested against EEG-weighted sleep scoring of selected scan epochs using receiver operating curve (ROC) analysis. Here, we compared EEG-verified awaking epochs (n = 30, EEG-weighted sleep score = 0) against sleep (n = 30, EEG-weighted sleep score ≥ 10). The ROC analysis of SE_ROI_ separated sleep from waking state with high accuracy (AUC=0.88, p<0.001, Fig.2f). Whole brain SE_GLOBAL_ accuracy had lower sensitivity for this discrimination (AUC=0.77, p<0.001, Supplementary Fig. 2c) compared to the occipital SE_ROI_ approach.

### Spatial MREG_BOLD_ SE change in vigilance and sleep

During routine fMRI scanning, decreasing vigilance during the first 3-5 minutes manifests in typical alterations in the BOLD signal (Tagliazucchi & Laufs, 2014). To analyze how decreased vigilance alters the SE of MREG_BOLD_ data, we compared the first five-minutes of scan 1 (A1 epoch 0-5 min) to the last five-minutes (A2 epoch, 5-10 min, i.e. A1 vs. A2 epochs). The SE in the final five-minutes of the MREG_BOLD_ signal was significantly lower than in the first five-minutes (p < 0.05, df 20, Fig 3b), although EEG sleep scoring did not detect significant sleep episodes on in either A1 or A2, each of which showed a single 30 sec N1 sleep episode. Eye closure (EC) has also been shown to reduce flow and increase BOLD fluctuations in visual cortex (Zou et al., 2015). In our study, the EC increased VLF power (as previously reported by (Zou et al., 2015), and indeed produced an even more pronounced SE decrease over the visual cortex and other posterior brain areas (A1 vs. EC, p < 0.05 FWE corrected, df 20, c.f. Fig 3b). The EEG sleep verification also showed higher prevalence of 30 sec time segments of sleep (in total 42 sleep time segments from 7/12 subjects) during awake eye closure. However, there was no difference in SE when comparing the effect of eye closure to scan duration (A2 vs. EC). The sleep epoch S2 showed the largest spatial extent with decreased in SE of the MREG_BOLD_ signal compared to awake epoch A1, covering most of the posterior brain area, Fig. 3a (A1 > S2, p < 0.05, df 20).

**Figure 3.**
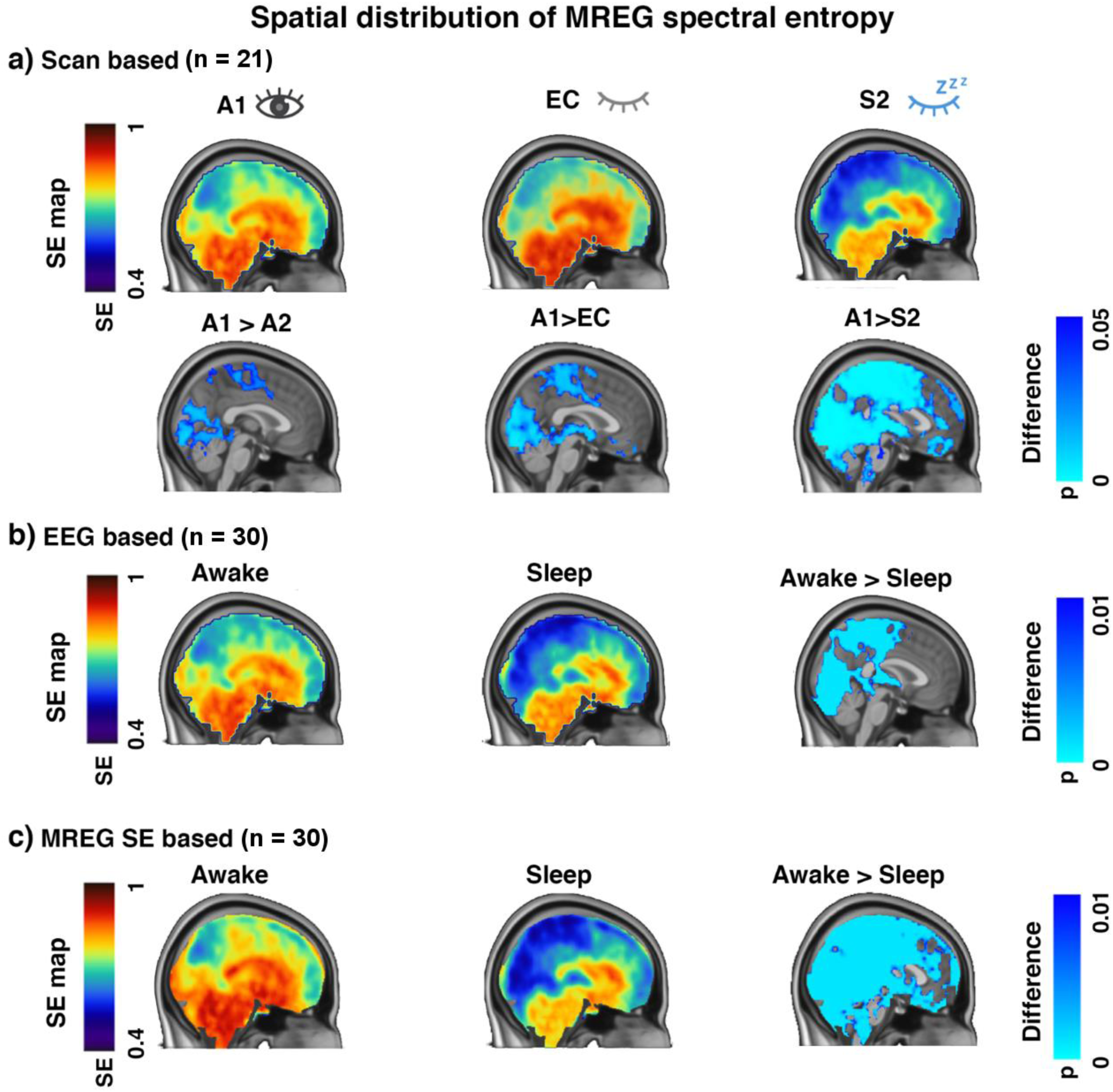
Spatial distribution of the sleep-induced reduction of full band MREG_BOLD_ spectral entropy (SE) signal in posterior and temporal parts of the brain. a) Scan based analyses illustrate (p < 0.05, df 20) how SE reduces in association with declining vigilance level with increasing scan duration (A1 (0-5 min) > A2 (5-10 min), eye closure (A1 > EC) and sleep scan effect (A1 > S2), when pooling data from all 21 subjects. When considering only awake (n = 30) and deepest sleeping scan epochs (n = 30), based on b) EEG-weighted sleep score and c) especially for the MREG_BOLD_ SE_ROI_ sleep score, the SE reductions had markedly higher statistical significance (p < 0.01, df 29). In practice, the SE changes occur in posterior brain areas, notably around primary sensory regions and their associative areas (visual, sensorimotor, auditory).

The EEG sleep scores also indicated that not all subjects were able to sleep during the morning multimodal fMRI scanning procedure, despite their sleep deprivation in the preceding night (Supplementary table 1-2). Therefore, to analyze *only* sleep effects with the clearest possible differentiation between waking and sleep states, we contrasted the 30 deepest individual sleep five-minute scan epochs with the 30 most awake scan epochs, based both on i) EEG-weighted sleep scores, and ii) MREG_BOLD_ SE_ROI_ criterion, in separate analyses. Irrespective of the criterion, the entire occipital, parietal and temporal lobes and cerebellum exhibited significant reductions in SE (Fig. 3 b-c, p < 0.01, df 29), whereas parts of the frontal lobe did not show SE changes. The EEG-scored and MREG_BOLD_ results showed overlapping SE reductions in the parietal and occipital regions, mostly concentrated near the somatosensory regions. However, the strength and spatial spread of the SE reduction were smaller in EEG-verified sleep scans compared to MREG_BOLD_ SE_ROI_-based results.

### Sleep increases the power of the brain pulsations

We further measured the power of each physiological brain pulsation with fast Fourier transform (FFT) analysis. Figure 4 shows the FFT power spectra changes from a ROI in the visual cortex (SE_ROI_, MNI coordinates: 12, -63, -6) and Figures 4 and 5 show the power changes across all voxels to evaluate the relative contributions of the physiological brain pulsations (VLF 0.008 - 0.1 Hz, respiratory 0.11 - 0.44 Hz, and cardiac 0.52 – 1.6 Hz) at different states of consciousness (Figs. 4-5). The VLF power was significantly higher already after eye closure during an EC scan (p < 0.05, df 20), and increased in the deepest average sleep scan S_2_ (p < 0.05, df 20) in comparison to the awake A1 scan in all 21 subjects. In EEG-verified sleep epochs, and more strongly in MREG_BOLD_ SE based sleep epochs, the VLF power increased (p < 0.05, df 29) in comparison to wakefulness (Figs. 4-5, Supplementary table 3). The power of respiration pulsations was similarly increased in sleep versus awake time segments (p < 0.05, df 29). The cardiac pulsation power increased to a lesser degree in both the EEG-based and SE_ROI_-based MREG data (p < 0.05, df 29). Previous work showed that heart rate variability increases in sleep, especially when respiratory frequencies fall below 0.5 Hz (Cajochen, Pischke, Aeschbach, & Borbély, 1994; Elsenbruch, Harnish, & Orr, 1999; Kondo et al., 2014). Similarly, our spectral analyses of the variability of the cardiac pulse frequency also increased from a single 1 Hz peak multiple frequency peaks ranging from 0.7 – 1.4 Hz (Fig. 4).

**Figure 4.**
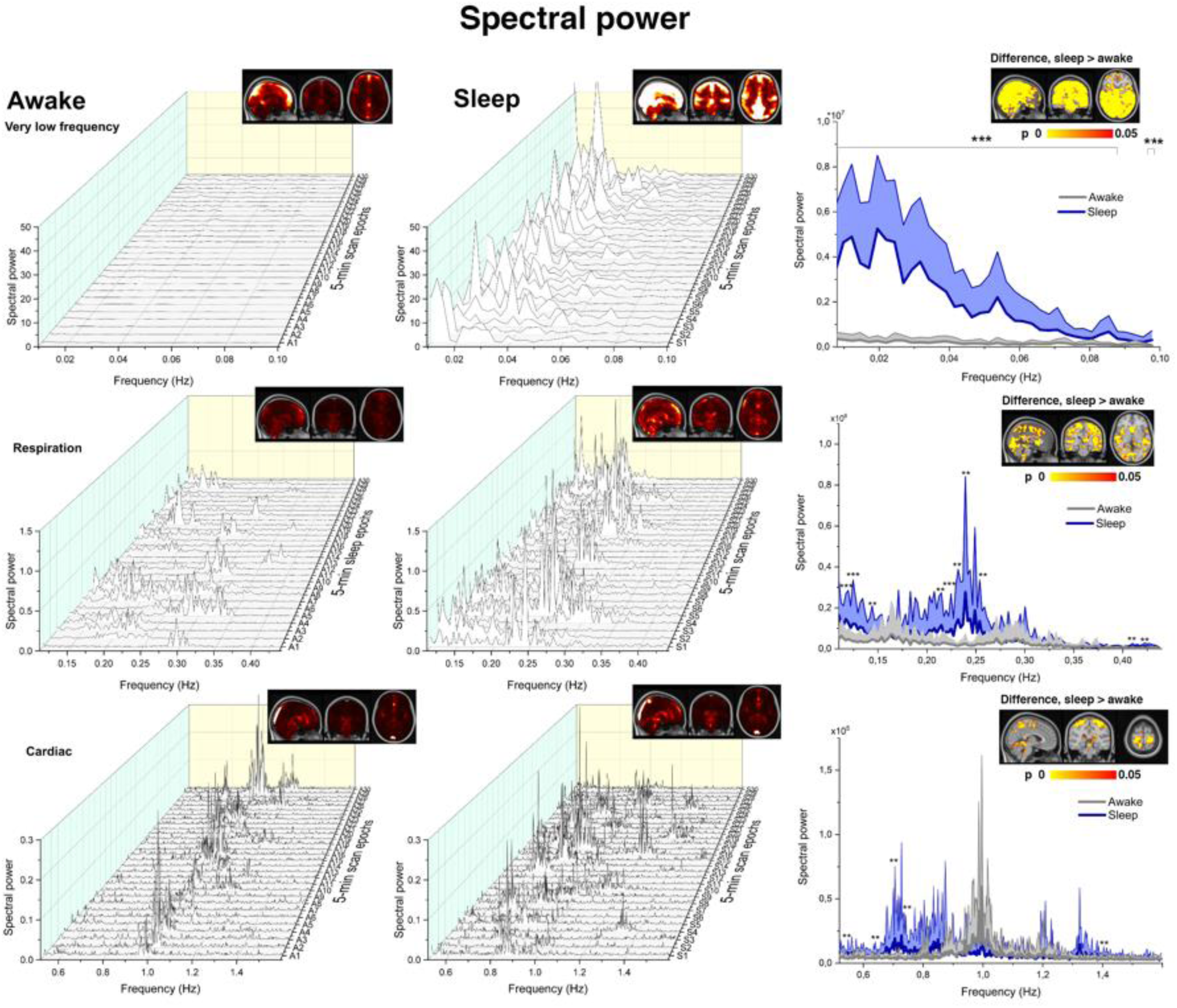
Sleep sharply changes the power spectrum of physiological brain pulsations. Power spectra from MREG_BOLD_ data from the visual cortex (MNI 12, -63, -6) from the most awake and deepest sleep data, as verified by SE_ROI_. The VLF pulsation is low in awake data, and increases markedly in sleep. The respiratory pulsation also increases significantly upon transition to sleep, although to a lesser extent than VLF. The cardiac pulsation power also increases and the variability of the cardiac pulsation power increases from a single 1 Hz peak into a range of frequencies from 0.7-1.4 Hz. (p < 0.001 ***, p < 0,01 **, p < 0,05 *, df 29).

**Figure 5.**
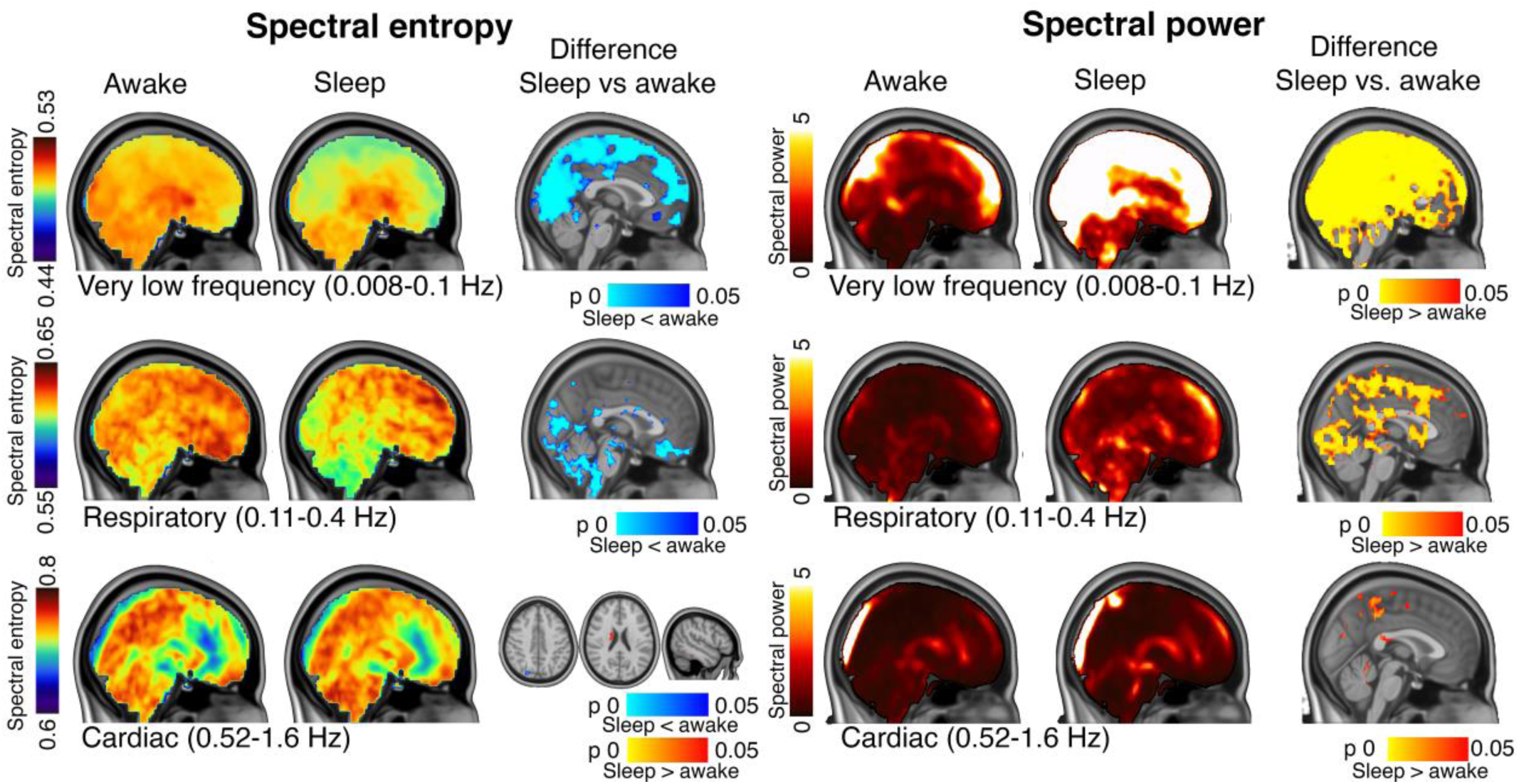
Spatial distribution of spectral brain pulsation entropy (SE) and power are linked to frequency band. Spectral entropy (left panel) and spectral power (right panel)in each pulsations frequency band in five-minute epochs of awake and deepest sleep based on MREG_BOLD_ SE_ROI_, and their statistical difference maps (n= 30 vs. 30 epochs, df 29). The three rows show results in VLF, respiratory and cardiac frequencies, respectively. Each frequency range shows significant increases in power of the physiological brain pulsations upon transition to sleep, although the spatial extents of the power and SE changes decline with increasing frequency band. The spatial overlap of the power and entropy increases also dimishes as a function of frequency, with largest overlap in the VLF range, partial overlap in the respiratory band, and no overlap in cardiac frequencies.

### Spatial extent of sleep induced entropy and power changes in each type of physiological brain pulsation

The spatial extent of the effect of sleep state on physiological pulsation entropy and power were also evaluated for each of the three pulsation frequency ranges, based both on MREG_BOLD_ SE_ROI_ and EEG-weighted sleep score data. Much as with SE_ROI_ sleep and vigilance results, the MREG_BOLD_ SE-based scoring also yielded robust FFT power spectrum results for each pulsation type. The largest power change was the increase in VLF power covering almost the entire posterior part of the brain, matching the pattern of SE change (MREG_BOLD_ SE vs. power *fslcc* = 0.9, Figs. 4-5). Furthermore, in solely awake data, the VLF power was significantly lower in the MREG_BOLD_ signal when verified based on MREG SE_ROI_ vs. EEG-verified (p < 0.001, df 29), indicating higher sensitivity of MREG_BOLD_ SE-based scoring also for detecting vigilance-related VLF changes (Supplementary Fig 3).

The respiratory pulsation power in the brain was also increased significantly in sleep (p < 0.01, df 29, Figs. 4-5). The peak mean change was located in upper posterior brain areas overlapping with the regions showing a VLF increase. Although clearly statistically significant, the increase was not as strong as the VLF power change (Figs. 4-5). The most significant power increase occurred along the bilateral central sulcus, thus involving both sensorimotor cortices, close to auditory cortices mainly of the left (dominant) hemisphere, and, over visual V1-V2 and V5 areas predominantly on the right side, and upper dorsal regions of the cerebellum. Again, the EEG-verified sleep data had smaller respiratory power increases than that from MREG_BOLD_ SE_ROI_ based, but both findings of increased power overlapped in the visual upper cortex (Supplementary Fig. 3). In contrast to the occipital VLF increase, the increase in respiratory frequency SE was located more towards the basal brain structures, with less overlap with the power changes in upper posterior regions.

The power of cardiac pulsation increased during sleep around sensorimotor areas and in the superior cerebellum (p < 0.05, df 29). Notably, the SE of the cardiac pulsation also showed a small drop in the parietal area, and interestingly two areas of *increased* SE, which stood in contrast to findings for VLF and respiratory frequencies. Eye closure EC (A1 vs. EC, p < 0.05, df 20) caused an increase in the power for cardiac and VLF entropy frequencies, but not in respiratory pulsations (Supplementary Fig.5).

## Discussion

We show in this study the spectral power of all three physiological pulsations of brain tissue increase during episodes of reduced vigilance and especially in the transition of sleep. Once falling asleep, there is a redistribution of the brain pulsations spectrum towards slower physiological brain pulsations, accompanied by a reduction in entropy in the spectral distribution. In fact, the SE of the brain pulsation signal in the visual cortex was an accurate marker for EEG-verified sleep epochs (ROC AUC=0.88, Fig 2). This suggests that SE calculated directly from MREG_BOLD_ brain scans serves for reliable data-driven monitoring of vigilance.

After separately analyzing the three main physiological pulsation in the ultrafast MREG_BOLD_ data, it became evident that the power increases in sleep with an inverse relation to the frequency. Thus, VLF fluctuations showed the strongest increase, followed by the faster respiratory and then cardiovascular pulsations. The respiratory pulsations increased only during verified sleep epochs, whereas the spatial extent of VLF and the cardiovascular pulsations increased even during brief lapses in awake vigilance, and certainly in the early transition to sleep following 24 hours of sleep deprivation (Figs. 4-5 & Supplementary Fig. 5). Even slight reductions in awake vigilance state were associated with significantly reduced the SE and increased the power of brain pulsations to ultrafast MREG_BOLD_. Because these waking vigilance changes did not qualify as sleep episodes in EEG when using standard 30 s epochs, we argue that physiological brain pulsations offer a more sensitive tool for detecting lapses in physiological vigilance in the transition to sleep.

### VLF brain pulsations increase during transient drops in vigilance and predominate in sleep

This is the first study to demonstrate brain-wide changes in the three major pulsation mechanisms in human brain during sleep. Our study was enabled by the ability of ultrafast MREG_BOLD_ with 10 Hz sampling rate to critically separate cardiorespiratory and VLF signals in human brain without aliased mixing of the faster activity over the VLF signal (Huotari et al., 2019). Earlier literature on the conventional BOLD signal showed increased VLF fluctuations in posterior brain regions during light sleep and episodes of low vigilance, which is in accord with present results showing markedly increased VLF power in sleep (Chang et al., 2016; Liu et al., 2018; Wen & Liu, 2016). Although the low frequency BOLD fluctuations are widely attributed to functionally connected hemodynamic oscillations that are tightly coupled with waking state neuronal activity (Hutchison et al., 2013; Korhonen et al., 2014), part of the low frequency activity has also been linked to vasomotor waves or slow sinusoidal hemodynamic oscillations at ∼0.1 Hz (Biswal, Bharat, Zerrin Yetkin, Haughton, & Hyde, 1995; Kiviniemi et al., 2000; Kiviniemi et al., 2016; Rayshubskiy et al., 2014; Wang et al., 2008).

Light sedation (Ramsey score 3) with intravenous midazolam increases brain VLF power significantly relative to the waking state (J. Kiviniemi et al., 2005). In this study, we show similarly increased power for very low pulsation frequencies (< 0.1 Hz) in conjunction with reduced SE of the pulsation. Hillmann and colleagues have similarly detected two separate VLF phenomena during anesthesia in mice; a 0.04 Hz fluctuation present only in anesthesia data overshadowed the faster hemodynamic changes coupled to spontaneous neuronal activity (Ma et al., 2016). Only upon removing the 0.04 Hz vasomotor fluctuation by filtering, there emerged the underlying the neuronal activity-coupled and functionally connected hemodynamic signal typical of spontaneous resting state networks. Furthermore, a recent paper showed a strong 0.05 Hz CSF pulsation in the fourth ventricle in human NREM sleep, which was coupled both the EEG theta power and 0.1 % BOLD signal oscillations (Fultz et al., 2019).

It seems plausible that, especially during sleep, the VLF BOLD signal might be affected by non-neuronal vasomotor waves at least to some degree.

Vasomotor waves are slow, seemingly spontaneous undulations in the arteriolar wall tension that control vessel wall pulsatility and local flow resistance, which consequently influence perfusion in downstream vascular territories (Preiss & Polosa, 1974). The vessel wall pulsatility in response to cardiac pressure waves serves as a driving force for the paravascular glymphatic CSF convection (Mestre et al., 2018). The force of the driving pulsation is dependent on various factors, including the circulatory perfusion pressure and elasticity of the vessel wall, both of which are under regulation from smooth muscle tonus in the arterial wall (Hadaczek et al., 2006). Vasomotor wave amplitude increases when blood perfusion pressure declines (Biswal, Bharat B. & Kannurpatti, 2009; Fujii, Heistad, & Faraci, 1990), and conversely, any increase in blood pressure reduces both the amplitude of vessel wall pulsation and paravascular CSF convection (Mestre et al., 2018). It seems likely that the increasing vasomotor waves in the arterial wall tension during sleep could facilitate vessel wall pulsatility and thereby promote paravascular CSF transport. If so, vasomotor waves during sleep could contribute to the glymphatic brain CSF transport and brain metabolite removal (Kiviniemi et al., 2016). The variations in arterial wall tension acutely affect blood vessel muscular thickness, and thus might modulate both perivascular space CSF volume and convection, but also contribute to parallel trafficking of water and metabolites via the *limitans externa* of the blood brain barrier wall into interstitium. This concept returns us to the theme that various pulsations are permissive to glymphatic flow.

### Respiratory pulsations increase specifically in sleep

For the first time we present evidence that sleep specific brain pulsations change in phase with the respiratory frequency. Although it is known that the physiological pulsation can affect the BOLD signal, this interaction has not been widely studied due to technical limitations of conventional fMRI (Huotari et al., 2019; Kiviniemi et al., 2016). The acquisition of ultrafast MREG_BOLD_ data enables robust separation of the cardiac and respiratory pulsations due to absence of signal aliasing (Huotari et al., 2019; Kiviniemi et al., 2016; Raitamaa et al., 2018) over the VLF frequencies.

During inspiration, the reduced intrathoracic pressure induces an outflow of venous blood from brain that is counterbalanced by an inward movement of CSF, as is a necessary consequence of the confinement of brain within the incompressible dural venous sinuses and intracranial space (Dreha-Kulaczewski et al., 2015; Dreha-Kulaczewski et al., 2017; Klose, Strik, Kiefer, & Grodd, 2000; Vinje et al., 2019; Yamada et al., 2013). This mechanism is bound to also effect the perivenous CSF space, which together with counter phase venous blood volume changes creates a perivenous CSF pump. This pump can function to transmit respiratory pulsations into the neuropil. In addition, the pulmonary ventilation and upper airway resistance increase up to four-fold during sleep (Sowho, Amatoury, Kirkness, & Patil, 2014; Trinder, Whitworth, Kay, & Wilkin, 1992; Wiegand, Zwillich, & White, 1989; Worsnop, Kay, Pierce, Kim, & Trinder, 1998). The altered intrathoracic ventilation pressures may consequently increase the driving (peri)venous pumping action in the brain, manifesting in altered respiratory frequency power in the MREG_BOLD_ signal (Dreha-Kulaczewski et al., 2015; Matsumae et al., 2019; Vinje et al., 2019).

The increases in respiratory and VLF brain pulsations with transition to sleep overlap in the posterior brain regions, and may be mediated by shared autonomic pressure control mechanisms. In contrast to VLF and cardiovascular brain pulsations, the respiratory pulsations do not increase with lower awake state vigilance (Supplementary Fig.5). Since respiratory brain pulsations increase during deeper stages of sleep than those required for increased cardiovascular pulsations, we hypothesize that the respiratory pulsations may be a main driving mechanism of glymphatic brain clearance in perivenous spaces, which further facilitates CSF convention during deep sleep. This prediction could be tested in future MREG_BOLD_ conducted across the entire sleep cycle. The observed discrepancy in the anatomical distributions of entropy and power changes of the respiratory pulsations is another matter for further research, possibly in relation to differential control of vascular mechanisms or pressure/flow characteristics of these anatomical areas.

### Cardiovascular brain pulsation change in sleep

The entropy and power of the cardiovascular pulsations do not range as widely as those of VLF and respiratory pulsations. The variability of the pulsation increases with sleep, in accordance with previous knowledge about sleep-related cardiovascular pulsation changes (Cajochen et al., 1994; Elsenbruch et al., 1999; Kondo et al., 2014). The increased cardiac variability during sleep has been linked in part to the respiratory frequency increase (Elsenbruch et al., 1999) and the various cardiorespiratory pulsations could well interact, given their closely connected intrathoracic physiology. Furthermore, the frequency distribution of brain cardiovascular pulsation widens and moves towards slow frequencies during sleep (Fig. 4, Supplementary table 4). This could reflect the increased and more efficient CSF convection that occurs along with reduced heart rate during sleep, resembling the anesthetized state (Hablitz et al., 2019). Drift in cardiovascular pulsation frequency and variability during sleep is most likely related to reduced sympathetic drive (Somers, Dyken, Mark, & Abboud, 1993), as are likewise the vasomotor wave amplitude increase and frequency reductions (Preiss & Polosa, 1974).

### Sleep physiology vs. MREG_BOLD_ spectral entropy

Previous studies have revealed episodes of EEG-verified (Tagliazucchi & Laufs, 2014) and self-reported sleep (Soehner et al., 2019) during “awake” resting state fMRI scanning conditions. Tagliazucchi et al. found that 30% of subjects fell asleep within three minutes of starting a resting state fMRI scanning (Tagliazucchi & Laufs, 2014), which is in accord with our present findings. We also show that SE of the MREG_BOLD_ can similarly serve as a metric for brain vigilance state. The 5-10 min awake scan epoch (A2) and eye closure (EC) interval both showed reduced SE and increased VLF brain pulsation compared to the initial 0-5 minute scan epoch (A1) (Figs. 3-5), see also (Supplementary table 3 & Fig. 5). Intriguingly, the A2 data showed reduced SE and increased VLF power compared to A1, without onset of EEG-verified sleep. The data also suggest that physiological brain pulsation changes may precede detectable neuronal activity changes, both during vigilance shifts and in sleep-wake transitions.

The cumulative EEG-based sleep scores based on AASM criteria had a linear correlation with visual cortex MREG_BOLD_ spectral entropy, c.f. Fig 2. The individual subject scan data and sleep scored epoch data both indicated a further decrease in the SE of the MREG_BOLD_ signal upon verified sleep onset. Similar to EEG entropy scores during anesthesia, the MREG_BOLD_ data possess the necessary spectral resolution for accurately determining sleeping state (Burioka et al., 2005; Liang et al., 2012; Mahon et al., 2008; Rodríguez-Sotelo et al., 2014; Vakkuri et al., 2005). Moreover, mapping of the MREG_BOLD_ SE can help localize the source of EEG brain oscillation changes in deep structures and across the whole brain.

Overall, the MREG_BOLD_ spectral power changes during sleep seem to be more widely distributed and spatially uniform than are the corresponding SE changes with respect to each pulsation mechanism. The spatial similarity of the power vs. entropy maps declines as a function of decreasing signal frequency. Thus, the map of VLF brain pulsation power increase closely matches the SE drop in the upper/posterior brain regions and is nearly identical to full band SE maps. The respiratory pulsations tend to increase in superior brain regions, while the entropy drops in the more basal structures. For the cardiac pulsation band, the power and entropy changes have little spatial overlap, for reasons yet to be explained.

## Conclusions

Sleep increases the power of all three physiological brain pulsations and reduces their complexity, as marked by decreased spectral entropy. The increased pulsation follow the rank order vasomotor > respiratory > cardiac pulsations, with declining spatial extents. The SE of brain pulsations declines over posterior brain areas even during transient drops in waking vigilance, making it a very sensitive indicator of physiological pulsation changes in the transition from awake to sleep states. Our results suggest that all three physiological brain pulsation mechanisms contribute in varying degrees to the increased glymphatic activity during sleep and may relate to the link between cardiovascular disease and Alzheimer’s disease in relation to impaired amyloid clearance.

## Material and Methods

### Subjects

Twenty-five subjects (aged 28.0 ± 5.9 years, 11 females) participated in the study, which was approved by the Regional Ethics Committee of the Northern Ostrobothnia Hospital District. Written informed consent was obtained from all participants, according to requirements of the Declaration of Helsinki. All subjects were healthy and met the following inclusion criteria: no continuous medication, no neurological nor cardio-respiratory diseases, non-smokers and no pregnancy. The subjects participated in two sessions, an Awake scan session in the afternoon (7.8 ± 1.2 hours sleep in previous night) and the Sleep scan session in the early morning following a sleepless night (Fig. 2a). The subjects were instructed not to consume caffeine during the four hours before the Awake scan session and eight hours before the Sleep scan session. Alcohol consumption was prohibited during the night before the scan.

### Data collection

All subjects were scanned in Oulu (Finland) using a Siemens MAGNETOM Skyra 3T (Siemens Healthineers AG, Erlangen, Germany) scanner with a 32-channel head coil. The subjects were scanned with ultrafast fMRI sequence, MREG, in synchrony with a previously described multimodal scanning setup (Korhonen et al., 2014). MREG is a single-shot sequence that undersamples k-space with an in/out stack-of-spiral trajectories in three dimensions (Assländer et al., 2013; Lee, Zahneisen, Hugger, LeVan, & Hennig, 2013; Zahneisen et al., 2012). The following parameters were used for MREG: repetition time (TR = 100ms), echo time (TE = 36ms), and flip angle (FA = 5°), field of view (FOV = 192 mm^3^) and 3 mm cubic voxel. Parameters for three-dimensional structural T1 MPRAGE were TR = 1900 msec, TE = 2.49 msec, FA = 9°, FOV = 240 mm^3^, and slice thickness 0.9 mm. MREG data were reconstructed using L2-Tikhonov regularization with lambda 0.1, with the latter regularization parameter determined by the L-curve method with a MATLAB recon-tool from sequence developers (Hugger et al., 2011).

During the Awake scan session, two MREG scans were recorded: a) ten-minute resting state, eyes open, awake scan (A_1-2_) fixating on a cross on the screen, a) five-minute resting state with EEG (EC) (Fig.2a). During the Sleep scan session three days later, one or two MREG sequences were recorded : a) the first ten-minute sleep scan (S_1-2_), b) the second ten-minute Sleep scan (S_3-4_), whereupon subjects were allowed to fall asleep *ad libitum*. They were advised to contact the staff if they were not feeling at all somnolent after the first scan, which would result in termination of the session. Due to individual differences in awake vigilance and in the ability to sleep in the MRI (despite presenting a dark and relatively quiet environment), the ten-minute scans were cut into five-minute segments (A_1_, A_2_, S_1_, S_2_, S_3_ and S_4_) to separate more effectively awake and sleep states for further analysis. An anatomical MR scan was performed at the end of both sessions.

EEG (0.01 Hz high-pass filtering) was recorded using Electrical Geodesics (EGI, Magstim Company Ltd, Whitland, UK) MR-compatible GES 400 system, with a 256-channel high density net. Electrode impedances were <50 kΩ and sampling rate 1 kHz (six subjects 250 Hz). Signal quality was tested outside the scanner room by recording 30?seconds of EEG with eyes open and eyes closed. Respiratory belt and fingertip peripheral SpO_2_ and anesthesia monitor data (ECG, fingertip peripheral SpO_2_ and end-tidal carbon dioxide (EtCO_2_), Datex-Ohmeda S/5 Collect software) were measured in synchrony with EEG, as described previously (Korhonen et al., 2014).

Two subjects were excluded because of previously un-known sleep apneic breathing patterns during the sleep scan and one because of low sleep score (only three hours of sleep) before the Awake scan session, compounded by excessive head motion during MREG acquisition. One subject was excluded from sleep analysis because of compromised MREG sleep data but was included in the awake vs. EC comparison (Supplementary table 1). One subject was included in the awake vs. sleep comparison, but no EC data were available.

### Smart-ring activity data

The subject sleep/wake status was monitored with the Oura-ring sleep tracker (www.ouraring.com) several days prior to scanning, data from the 24 hours preceding both scanning sessions was further analyzed. The smart ring records upper limb motions (3-D accelerometer, 50 Hz), photoplethysmogram (250 Hz) and skin temperature (1/min). The ring had 96% sensitivity to detect sleep, as previously documented in an independent validation study among healthy young sleepers (de Zambotti, Rosas, Colrain, & Baker, 2019). In this study, data from the ring was used to confirm that subjects remained awake during the night before the Sleep scan session. Ring classification of sleep is done in 30 s epochs. To initiate a sleep period, two minutes of full rest is required, where full rest is detected if the band pass filtered hand acceleration signals stay consistently below 64 mGs in all three dimensions. Additionally, ring accelerometer data must indicate low activity levels and the photoplethysmogram signal must indicate stable heart rate dynamics and pulse amplitude characteristics, if sleep is to be registered. Oura ring data were available for 21 subjects. Total sleep duration was calculated from the 24 hour periods preceding both scanning sessions (Supplementary Fig. 1).

### CANTAB tests

At the beginning of scanning sessions, prior to actual scans, the subjects underwent a reaction time (RTI) and paired-associate learning (PAL) tests from the CANTAB (Cambridge Neuropsychological Test Automated Battery, Cambridge, UK) test battery, performed on a tablet computer. The CANTAB test is widely used to study cognitive state and vigilance.

### Preprocessing and analysis of MREG data

After reconstruction, MREG data were preprocessed and analyzed using FSL (5.09 BET software (Jenkinson, Mark, Beckmann, Behrens, Woolrich, & Smith, 2012; Smith, 2002)), AFNI (Analysis of Functional NeuroImages, v2) and MATLAB (vR2018b; The Math Work, Natick, MA). The brain was extracted from structural 3D MPRAGE volumes with parameters f=0.25 and g=0.22 using neck clean-up and bias field correction options (Smith, 2002). The functional data preprocessing was done in the FSL pipeline. The data were high-pass filtered with a cut-off frequency of 0.008 Hz, and head motions were corrected with FSL 5.08 MCFLIRT software (Jenkinson, M., Bannister, Brady, & Smith, 2002). Spatial smoothing was done with *fslmaths* using a 5 mm FWHM Gaussian kernel. The highest spikes in the MREG data time series were removed using the *3dDespike* function in AFNI. Data were registered into MNI space at 3-mm resolution for comparable analysis between subjects and separated into five-minute scan epochs (A_1_-S_4_, 2861 time points) using FSL function *fslroi*.

### Preprocessing and analysis of EEG data

EEG recordings were preprocessed using the Brain Vision Analyzer (Version 2.1; Brain Products) after converting to acceptable format via BESA Research (Version 7.0). Gradient artifacts due to static and dynamic magnetic fields during MRI data acquisition and the ballistocardiographic (BCG) artifacts were corrected using the average artifact subtraction method (Allen, Polizzi, Krakow, Fish, & Lemieux, 1998; Allen, Josephs, & Turner, 2000). Checking for absence of gradient of BCG artifacts was done manually by visual inspection. After preprocessing, EEG data were visualized according to the 10-20 system instructions for sleep state scoring. Two experienced specialists in clinical neurophysiology, who were specially trained in sleep studies scored the EEG data. The scoring was performed in 30 s time segments following the AASM guidelines for clinical sleep studies (AASM, 2017), and the final sleep state scoring was obtained by consensus. Using established criteria, EEG epochs were scored as awake, N1 (light sleep), N2 (intermediate sleep with sleep spindles and/or K-complexes), N3 (slow wave sleep) or REM (sleep with rapid eye movements).

The number of 30 s time segments was multiplied with a sleep weight coefficient (awake=0, N1=1, N2=2, N3=3) for EEG based sleep scores for a given scan epoch. All 30 s time segments of a five-minute scan epoch were added to the EEG weighted sleep score. A fully awake five-minute session corresponds to a minimum EEG weighted sleep score of 0 and a five-minute epoch of N3 sleep corresponds to a maximum EEG weighted sleep score of 30. Data were included if sleep states and SE were both available (A_1_: n=12, A_2_: n=12, EC: n=13, S_1_: n=12, S_2_: n=12, S_3_: n=10 and S_4_: n=10, Supplementary table 2).

Although fMRI scanning introduces artifacts in the simultaneous EEG, we employed standard EEG clean-up strategies and obtained qualitatively adequate EEG for scoring from 12 subjects. EEG was measured for all subjects, but data from a subset of (n=10) subjects was inadequate for EEG sleep scoring due to a data cable artifact that was discovered only after final scanner artifact removal. However, the SE of MREG_BOLD_ proved to be sensitive tool to evaluate sleep induced physiological changes, with an accuracy for detecting sleep at least as good as that from EEG-scored sleep data. We did not undertake pre-screening for sleep apnea or other sleep disorders, since all included subjects were young, and reported themselves to be healthy. However, as noted above, breath interruptions during sleep scans were detected in two subjects, leading to exclusion of their data from further analysis.

### Vigilance estimation with MREG signal spectral entropy against EEG verification data

SE is a measure of spectral entropy of data and, as such, describes the amount of information contained with a given signal. The SE method is based on treating the normalized FFT power spectrum as a probability distribution and using this distribution to calculate the Shannon entropy (Shannon, 1948). Because of the 10 Hz sampling rate of MREG data, calculation of SE becomes a feasible tool to estimate vigilance changes and sleep from fMRI, thus presenting an alternative to EEG SE analysis (Kumar, Ramaswamy, & Nath Mallick, 2013; Mahon et al., 2008; Vakkuri et al., 2005).

SE entropy was calculated with MATLAB R2019 using the *pentropy* for global MREG signals, as used in earlier fMRI studies to measure arousal fluctuations (Chang et al., 2016; Liu et al., 2018) and also on a voxel-by-voxel basis. SE analyses were first performed on all five-minute MREG epochs. Sleep effects were analyzed from both EEG-verified awake (EEG-weighted sleep score = 0, n=30) and sleep states (EEG-weighted sleep score > 10, n=30) using the five-minute scan epochs that included over > 50% NREM N1-N3 sleep (Supplementary tables 1-2), and directly from fMRI data by applying the visual cortex ROI (SE_ROI_, MNI: x 12, y -63, z -6), MREG_BOLD_ SE_ROI_ verified awake (five-minute epochs with highest SE, n = 30) and sleep (five-minute epochs with lowest SE, n =30). Normal awake vigilance effects were analyzed from A_1_ vs. A_2_ (n=22), A_1_ vs. EC (n=21) and A_2_ vs. EC (n=21). The SE_ROI_ was also analyzed by linear regression analysis with EEG-based sleep weighted score. The SE_GLOBAL_ and SE_ROI_ values were used to verify awake (EEG-weighted sleep score = 0, n=30) and sleep status (EEG-weighted sleep score >10, n=30) against EEG based sleep score in receiver operating curve (ROC)-analysis for combined sensitivity and specificity of the SE MREG (Origin, 2019).

### FFT power spectral analysis of MREG signal

The VLF range of 0.008-0.1 Hz was chosen to get maximum coverage possible of the low frequencies without cross-talk with respiratory frequencies. Group cardiorespiratory frequency ranges were obtained from individual anesthesia and/or scanner physiological monitoring signals for FFT power analysis: respiratory (0.11-0.44 Hz) and cardiac (0.52-1.6 Hz). Each VLF, respiratory and cardiac range FFT power density map was calculated with the *3dPeriodogram* function in AFNI for A_1_, EC and S_1-2_ scans. Awake and sleep states were verified in two ways: 1) EEG verification, where awake state includes those five-minute scan epochs that have EEG-weighted sleep score = 0 (n=30) whereas sleep state includes those epochs that have EEG-weighted sleep score > 10 (n=30) and 2) MREG_BOLD_ SE_ROI_ verification, where awake state includes those five-minute scan epochs that have highest SE (n=30) and accordingly sleep state includes those epochs that have lowest SE (n=30). The five-minute epochs consisted of 2861 time points, and non-uniform FFT (NFFT) was conducted with 4096. Data were tapered using the Hamming window and zero-padded to the FFT length. The sum of VLF, respiratory and cardiac FFT power was calculated from global and voxel-wise MREG using *fslroi* (choosing the frequency range with bins that refer to frequency range). With FFT power analysis, we studied how brain pulsations are altered by both vigilance level and sleep state. VLF FFT power results were correlated with SE results using *fslcc* function to evaluate their relationship.

### Statistical information

We used FSL *randomise* with 10,000 permutations, TFCE (Threshold-Free Cluster Enhancement) and family-wise error (false positive) control along with a two-sample paired t-test model to compare different scan epochs (A1, A2, EC, S1, S2, S3, S4, df 20) and a two-sample unpaired t-test model to compare different states (awake and sleep, df 29) in the voxel-wise SE and the FFT power analysis. EEG-weighted sleep score was correlated with MREG_BOLD_ SE using linear regression (Pearson correlation, F-test). ROC curves were plotted to show accuracy of SE to separate sleep from awake state. For Oura-ring activity data, CANTAB performance, cardiorespiratory signals and head motion, statistical differences were calculated by two-sample paired t-test model using SPSS (IBM SPSS Statistics 26).

### Cardiorespiratory signals

SpO_2_ and EtCO_2_ signals were used to verify group cardiorespiratory frequency ranges based on individual ranges recorded for each subject. In case of poor quality or missing EtCO_2_ datasets, the respiration belt data were used to verify the frequency range. Frequency location of the maximum peak was determined and used to calculate the differences between A_1-2_, EC, S_1-2_ and S_3-4_.

### Motion

In addition to MCFLIRT correction and spike removal of data, we further excluded any effects of head motion. MCFLIRT relative motion signal was separated to five-minute time segments for comparison with other analysis, and the mean relative head motion (mm) was calculated. Values were demeaned and used as covariates in statistical analyses of voxel-wise fMRI.

## List of Abbreviations

AASM: American Academy of Sleep Medicine,
AFNI: analysis of functional neuroimaging software toolbox,
AUC: area under curve,
A1-A2: awake scan 1-2,
BCG: ballistocardiographic,
BOLD: blood oxygen level dependent,
CANTAB: Cambridge Neuropsychological Test Automated Battery,
CO_2_: carbon dioxide,
CSF: cerebrospinal fluid,
DC: direct current,
df: degrees of freedom,
EC: eyes closed scan,
EEG: electroencephalography,
EGI: Electrical Geodesics,
FA: flip angle,
FFT: fast Fourier transform,
fMRI: functional magnetic resonance imaging,
FMRIB: functional magnetic resonance imaging of the brain,
FOV: field of view,
FSL: FMRIB software library,
FWE: family wise error,
MATLAB: matrix laboratory software,
MCFLIRT: motion correction using FMRIB’s linear image registration tool,
MNI: Montreal Neurological Institute,
MPRAGE: magnetization prepared rapid gradient echo,
MREG: magnetic resonance encephalography,
light sleep: N1,
intermediate sleep with sleep spindles and/or K-complexes: N2,
slow wave sleep: N3,
NREM: non-rapid eye movement,
OFNI: Oulu functional neuroimaging group,
PAL: paired-associate learning,
sleep with rapid eye movements: REM,
ROC: receiver operating curve,
ROI: region of interest,
RTI: reaction time,
S1–S4: sleep scans 1-4,
SE: spectral entropy,
SpO_2_: fingertip oxygen partial saturation,
TE: echo time,
TFCE: threshold-free cluster enhancement,
TR: repetition time,
VLF 0.008 - 0.1 Hz: very low frequency.

## Acknowledgements

We would like to thank all study subjects for their participation in the study. We also thank Inglewood Biomedical Editing for proof-reading and language checking, Tuomas Konttajärvi for assistance in measurements and preprocessing of EEG data, Jani Häkli, Annastiina Kivipää, Tarja Holtinkoski, Aleksi Rasila, Taneli Hautaniemi, Miia Lampinen and others who assisted in measurements or participated otherwise. We are thankful for provided devices and data by Oura and Jussi Kantola making the reconstruction of MREG by the CSC – IT Center for Science Ltd., Finland.

## Author contribution statement

HH, VKo, SCH, JK, MJ, VB, MN, VKi designed research; HH, VKo, NH, MJ, VR, VB, VKi performed research; HH, VKo, JP, MK, TV, NH, LR, JT, JK, TT, VB, VKi analyzed the data, HK provided Oura measurement device and data, HH, VKo, SCH, JP, MK, TV, NH, LR, JT, JK, MJ, VR, TT, VB, HK, MN, VKi wrote the paper.

## Competing interests

Oura Health Ltd.’s provided the surveillance Oura rings for the study.

## Materials & Correspondence

Data of this study are available from the corresponding author (VKi) upon request.

## Funding

This work was supported by Uniogs/MRC Oulu DP-grant (HH, JK), Pohjois-Suomen Terveydenhuollon tukisäätiö (HH, VKo), JAES-Foundation (VKi), Academy of Finland and Aivosäätiö TERVA grant 314497 (VKi), Academy of Finland Grants 275342 (VKi), The SalWe Research Program for Mind and Body (Tekes—the Finnish Funding Agency for Technology and Innovation, Grant No. 1104/10) (VKi), Finnish Medical Foundation (VKi), Finnish Neurological Foundation, KEVO grants from Oulu University hospital (VKi), Epilepsy Research Foundation (JK), Finnish Cultural Foundation, North Ostrobothnia Regional Fund (JK), Orion Research Foundation sr (JK), Tauno Tönning Foundation (JK), The University of Oulu Scholarship Foundation (JK), Maire Taponen Foundation sr (JK), Finnish Brain Foundation sr (JK), Instrumentarium Science Foundation sr (JK)

