## Supplemental files for "Sleep-specific changes in physiological brain pulsations"

### Supplementary information:

##### Cardiorespiratory signals

Based on cardiorespiratory signals, respiration of all subjects was included in the 0.11-0.44 Hz band and cardiac cycle in 0.52-1.6 Hz, so those bands were used for further analysis. Additionally, we found that respiratory frequency was significantly higher in $A_{1-2}$ compared to $S_{1-2}$ (0.26 ± 0.06 vs. 0.24 ± 0.05 Hz, , p=0.006), and cardiac frequency was significantly higher in $A_{1-2}$ compared to EC (1.02 ± 0.14 vs. 0.98 ± 0.13Hz, p=0.00003), $S_{1-2}$ (1.02 ± 0.14 vs. 0.92 ± 0.12 Hz, p=0.006) and $S_{3-4}$ (0.99 ± 0.12 vs. 0.93 ± 0.12 Hz, p=0.03) (Supplementary table 4).

##### Motion

No difference in relative mean displacement (mm) was found in five-minute segments of $A_1$ except for $S_2$, when the mean relative motion was higher (0.0191 ± 0.007 mm) compared to $A_1$ (0.0183 ± 0.006 mm, p=0.014) (Supplementary Fig.6). The motion was used as a regressor in all spatial MREG map analyses in *randomise*.

**Supplementary table 1. Population of subjects and sample size for analysis.** The subject IDs are those used in other tables tables. $A_{1\text{-}2}$, awake eyes open (ten-minutes); EC, eyes closed (five-minutes); $S_{1\text{-}2}$, first sleep scan (ten-minute) following sleep deprivation, $S_{3\text{-}4,}$ second sleep scan (ten-minutes).

| | | | Awake | | Sleep | | |
|---|---|---|---|---|---|---|---|
| Subject ID N=22 | Oura n=21 | Cantab n=22 | $A_{1\text{-}2}$ n = 22 | EC n = 21 | $S_{1\text{-}2}$ n = 21 | $S_{3\text{-}4}$ n = 19 | EEG sleep state available n =12 |
| 1 | x | x | x | x | | | |
| 2 | x | x | x | x | x | x | |
| 3 | x | x | x | x | x | x | x |
| 4 | x | x | x | x | x | x | |
| 5 | x | x | x | x | x | x | x |
| 6 | x | x | x | x | x | x | x |
| 7 | x | x | x | x | x | x | x |
| 8 | x | x | x | x | x | x | x |
| 9 | x | x | x | x | x | x | |
| 10 | | x | x | x | x | | x |
| 11 | x | x | x | x | x | x | |
| 12 | x | x | x | x | x | x | x |
| 13 | x | x | x | x | x | x | |
| 14 | x | x | x | | x | x | x |
| 15 | x | x | x | x | x | x | x |
| 16 | x | x | x | x | x | x | |
| 17 | x | x | x | x | x | x | |
| 18 | x | x | x | x | x | x | x |
| 19 | x | x | x | x | x | | |
| 20 | x | x | x | x | x | x | x |
| 21 | x | x | x | x | x | x | |
| 22 | x | x | x | x | x | x | x |

**Supplementary table 2. The amount of NREM sleep was higher during the sleep scan session than in the awake scan session.** Mean ± standard deviation (%) amount of waking, N1-N3 and total sleep times were calculated from the number of sleep-scored EEG epochs in each condition. The highest amount of sleep was achieved in $S_2$ (mean across subjects 87%). Unknown epochs are due to artefact. A1 (awake 1 time segment, five min); A2 (awake 2 time segment, five min); EC (eyes closed, five min); S1 (sleep 1 time segment, five min); S2 (sleep 2 time segment, five min); S3 (sleep 3 time segment, five min); S4 (sleep 4 time segment, five min).

| | Awake scan session | | | Sleep scan session | | | |
|---|---|---|---|---|---|---|---|
| Mean ± STD (%) | $A_1$ n=12 | $A_2$ n=12 | EC n=13 | $S_1$ n=12 | $S_2$ n=12 | $S_3$ n=10 | $S_4$ n=10 |
| Wake | 99 ± 3 | 99 ± 3 | 68 ± 39 | 22 ± 19 | 11 ± 23 | 23 ± 32 | 25 ± 35 |
| N1 | 1 ± 3 | 1 ± 3 | 28 ± 39 | 47 ± 25 | 40 ± 33 | 40 ± 22 | 36 ± 30 |
| N2 | 0 | 0 | 4 ± 14 | 25 ± 19 | 46 ± 38 | 37 ± 29 | 37 ± 36 |
| N3 | 0 | 0 | 0 | 0 | 1 ± 3 | 0 | 0 |
| Unknown | 0 | 0 | 0 | 6 | 2 | 0 | 2 |
| Sleep mean (%) | 1 | 1 | 32 | 72 | **87** | 77 | 73 |

**Supplementary table 3. Strengths of pulsations measured by FFT power analysis of very low, respiratory and cardiac frequency were increased during sleep scans and EEG verified sleep compared awake.** Sum of global FFT power in very low (0.008-0.1 Hz), respiratory (0.11-0.44 Hz) and cardiac frequency (0.52-1.6 Hz). $A_1$, awake time segment eyes open, 5 min; EC, eyes closed five min, $S_{1\text{-}4}$, Sleep scan five-minute time segments. Mean ± standard deviation and p-value.

| | Awake vs. EC/sleep | **Sum of global MREG power (a.u. *10$^6$)** **Very-low frequency 0.008-0.1 Hz** | | | | | | | p-value |
|---|---|---|---|---|---|---|---|---|---|
| n=21 | $A_1$ vs. EC | 15.7 | ± | 13.1 | < | 21.1 | ± | 14.3 | **0.03*** |
| n=21 | $A_1$ vs. $S_1$ | 16.9 | ± | 13.3 | < | 34.0 | ± | 16.6 | **0.00001***** |
| n=21 | $A_1$ vs. $S_2$ | 16.9 | ± | 13.3 | < | 38.7 | ± | 21.7 | **0.0002***** |
| n=19 | $A_1$ vs. $S_3$ | 17.2 | ± | 13.8 | < | 29.7 | ± | 13.2 | **0.0003***** |
| n=19 | $A_1$ vs. $S_4$ | 17.2 | ± | 13.8 | < | 33.1 | ± | 17.3 | **0.0005**** |
| | | **Respiratory frequency 0.11-0.44 Hz** | | | | | | | |
| n=21 | $A_1$ vs. EC | 5.3 | ± | 2.7 | ~ | 6.3 | ± | 5.5 | 0.2871 |
| n=21 | $A_1$ vs. $S_1$ | 5.2 | ± | 2.6 | < | 8.2 | ± | 4.6 | **0.002**** |
| n=21 | $A_1$ vs. $S_2$ | 5.2 | ± | 2.6 | < | 10.5 | ± | 8.1 | **0.004**** |
| n=19 | $A_1$ vs. $S_3$ | 5.1 | ± | 2.6 | ~ | 6.8 | ± | 4.8 | 0.1039 |
| n=19 | $A_1$ vs. $S_4$ | 5.1 | ± | 2.6 | < | 7.6 | ± | 4.9 | **0.02*** |
| | | **Cardiac frequency 0.52-1.6 Hz** | | | | | | | |
| n=21 | $A_1$ vs. EC | 3.8 | ± | 0.8 | ~ | 3.9 | ± | 1.0 | 0.2576 |
| n=21 | $A_1$ vs. $S_1$ | 3.8 | ± | 0.9 | < | 4.5 | ± | 1.2 | **0.02*** |
| n=21 | $A_1$ vs. $S_2$ | 3.8 | ± | 0.9 | < | 4.4 | ± | 1.2 | **0.03*** |
| n=19 | $A_1$ vs. $S_3$ | 3.8 | ± | 0.8 | ~ | 4.2 | ± | 1.2 | 0.0574 |
| n=19 | $A_1$ vs. $S_4$ | 3.8 | ± | 0.8 | ~ | 4.3 | ± | 1.2 | 0.0552 |

**Supplementary table 4. Analysis of cardiorespiratory signals showed that cardiac frequency was decreased during eyes closed and sleep scans compared to awake, and respiratory frequency was decreased during sleep.** Maximum FFT peak location (Hz) from cardiorespiratory signals (respiration, $etCO_2$ and cardiac $SpO_2$). Mean ± standard deviation and p-value.

| | **Resp, $etCO_2$ (Hz)** | | | p-value | | **Card, $SpO_2$ (Hz)** | | | p-value |
|---|---|---|---|---|---|---|---|---|---|
| $A_{1-2}$ vs. EC (n=18) | 0.26 ± 0.06 | ~ | 0.24 ± 0.07 | 0.16 | n=18 | 1.02 ± 0.14 | > | 0.98 ± 0.13 | **0.00003*** ** |
| $A_{1-2}$ vs. $S_{1-2}$ (n=17) | 0.26 ± 0.06 | > | 0.24 ± 0.05 | **0.006** ** | n=18 | 1.02 ± 0.14 | > | 0.92 ± 0.12 | **0.0006*** ** |
| $A_{1-2}$ vs. $S_{3-4}$ (n=16) | 0.27 ± 0.07 | ~ | 0.25 ± 0.06 | 0.16 | n=15 | 0.99 ± 0.12 | > | 0.93 ± 0.12 | **0.03*** |
| EC vs. $S_{1-2}$ (n=18) | 0.24 ± 0.06 | ~ | 0.24 ± 0.05 | 0.81 | n=18 | 0.98 ± 0.13 | ~ | 0.94 ± 0.13 | 0.05 |
| EC vs. $S_{3-4}$ (n=16) | 0.25 ± 0.06 | ~ | 0.24 ± 0.06 | 0.95 | n=15 | 0.95 ± 0.12 | ~ | 0.94 ± 0.13 | 0.81 |

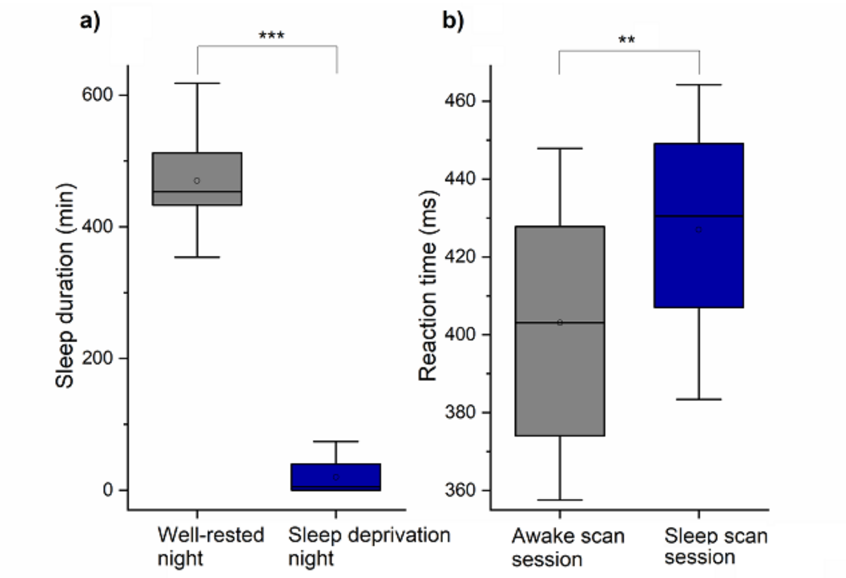


***Supplementary Figure 1. Sleep deprivation night with almost no sleep and lower reaction time after sleep deprivation predict low vigilance level during subsequent sleep scan session.*** *a) Sleep duration was significantly lower (***$p < 0.001$, df 20) in sleep deprivation than during the normal night prior to respective scans. b) Reaction time was significantly (**$p < 0.01$, df 21) longer after sleep deprivation night prior to Sleep scan session than prior to Awake scan session.*

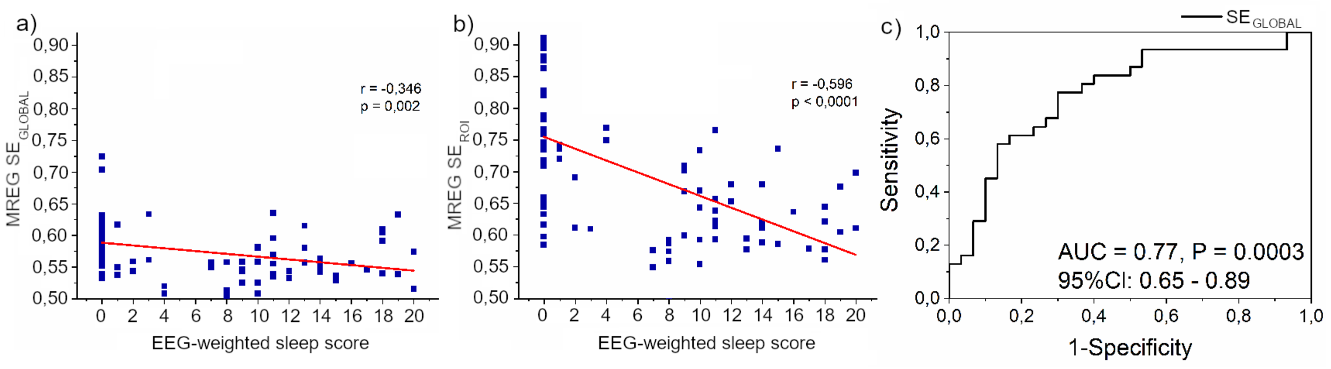


***Supplementary Figure 2. Spectral entropy of MREG and EEG-based brain signals reflect vigilance.*** *a) The MREG* $SE_{GLOBAL}$ *correlates with EEG-weighted sleep score (r = -0.35, p = 0.002, df 29). b) The MREG* $SE_{ROI}$ *correlates even better with the EEG-weighted sleep score (r –0.6, p < 0.001, df 29). c) ROC analysis shows that MREG* $SE_{GLOBAL}$ *served to separate sleep from awake state with a high accuracy (AUC=0.77, p < 0.001, df 29).*

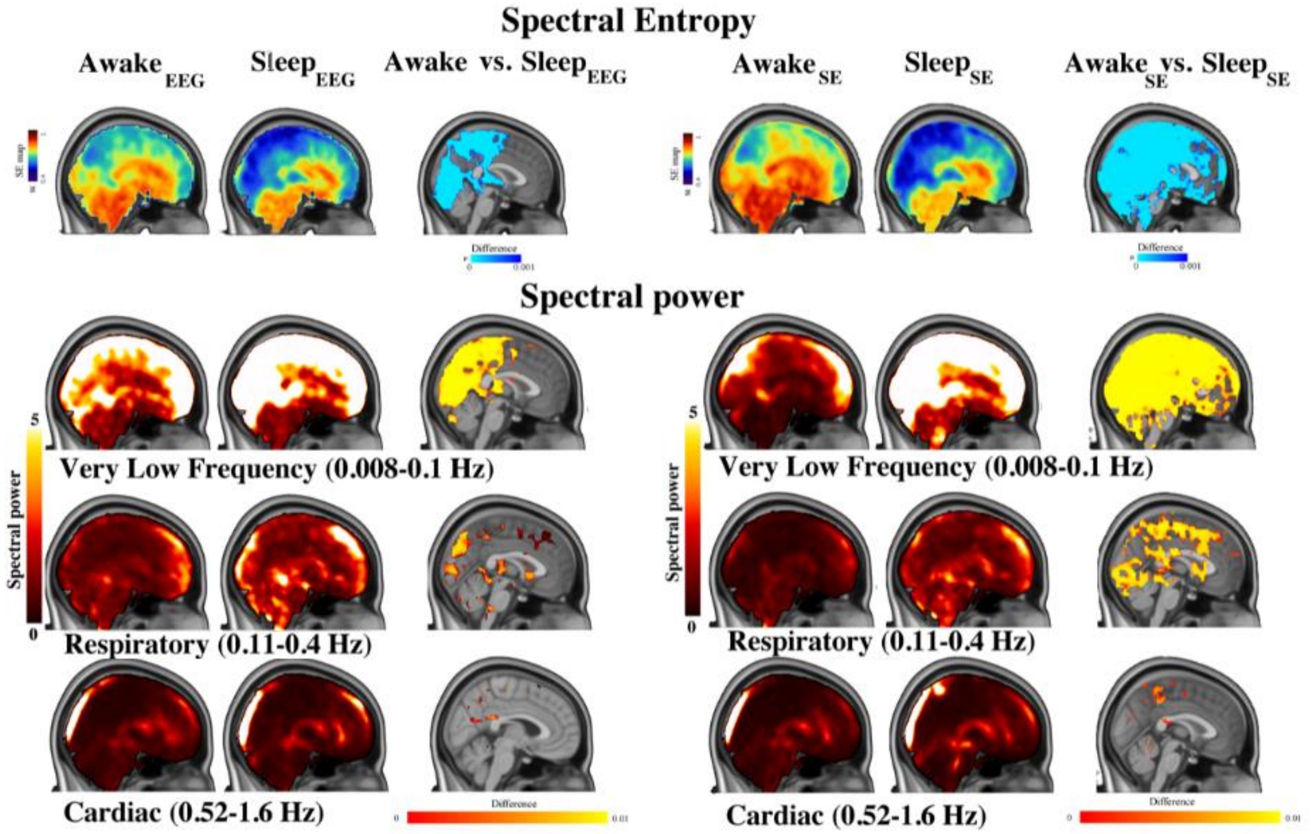


***Supplementary Figure 3. Spatial distribution of spectral brain pulsation entropy and power differences between EEG-verified (left panel) and MREG $SE_{ROI}$-verified (right panel) $MREG_{BOLD}$ data in the most awake and asleep states.*** *Spectral entropy (SE) in full band (top) and spectral power in each pulsation frequency (bottom) from most awake and deepest sleep five-minute scan epochs and their statistical difference based on both EEG-verified and MREG $SE_{ROI}$-verified $MREG_{BOLD}$ data (n = 30 vs. n= 30 epochs). The lower three rows depict spectral power results in VLF, respiratory and cardiac frequencies (from top to bottom). Spatial distribution of the sleep-induced reduction of SE signal in posterior parts of the brain. Each frequency range shows significant increases in power of the physiological brain pulsations after sleep, and the spatial extent of this changes declines as a function of increasing frequency. However, MREG $SE_{ROI}$-verified results are spatially more widespread compared to corresponding EEG-verified data both with respect to SE and spectral power.*

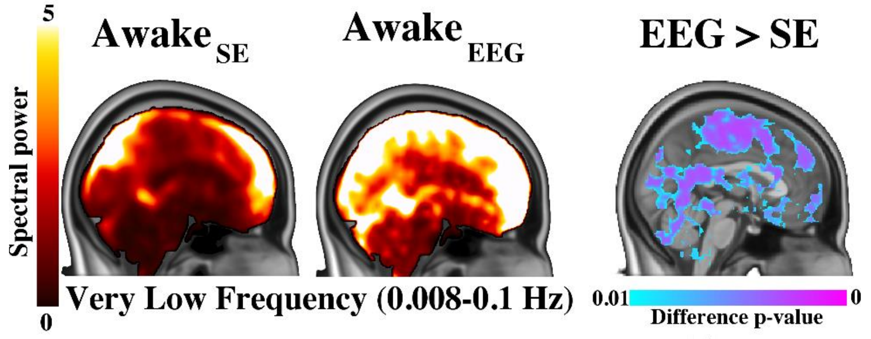


***Supplementary Figure 4. Spatial distribution of spectral brain pulsation power between MREG $SE_{ROI}$-verified awake (Awake$_{SE}$) and EEG-verified awake (Awake$_{EEG}$) MREG$_{BOLD}$ data in the very low frequency band and their difference map (EEG > SE).*** *Spectral power is markedly ($p < 0.01$, df 20) higher in EEG-verified awake image, which suggests that MREG $SE_{ROI}$ is a more sensitive indicator of awake status. Note that subjects are already partly sleeping in the Awake$_{EEG}$, although this cannot be discerned from EEG data scored based on AASM guidelines in 30 s epochs.*

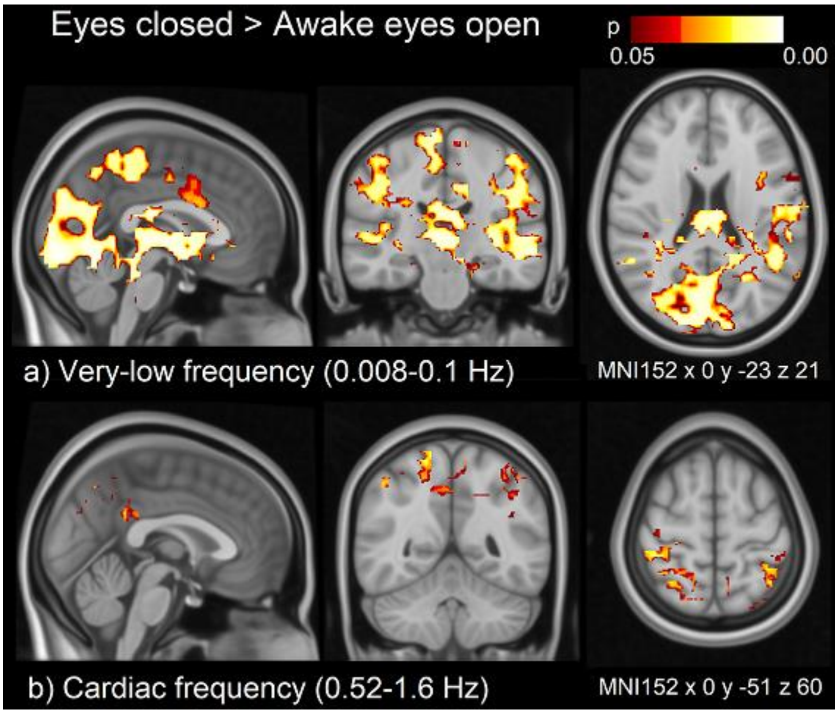


***Supplementary Figure 5. Spatial distribution of spectral brain pulsation power with eyes closed while awake.*** *Spectral power was significantly ($p<0.05$) higher during eyes closed compared to eyes open in a) very-low (0.008-0.1 Hz) and b) cardiac frequency (0.52-1.6 Hz) band, but no such difference was found for respiratory frequency.*

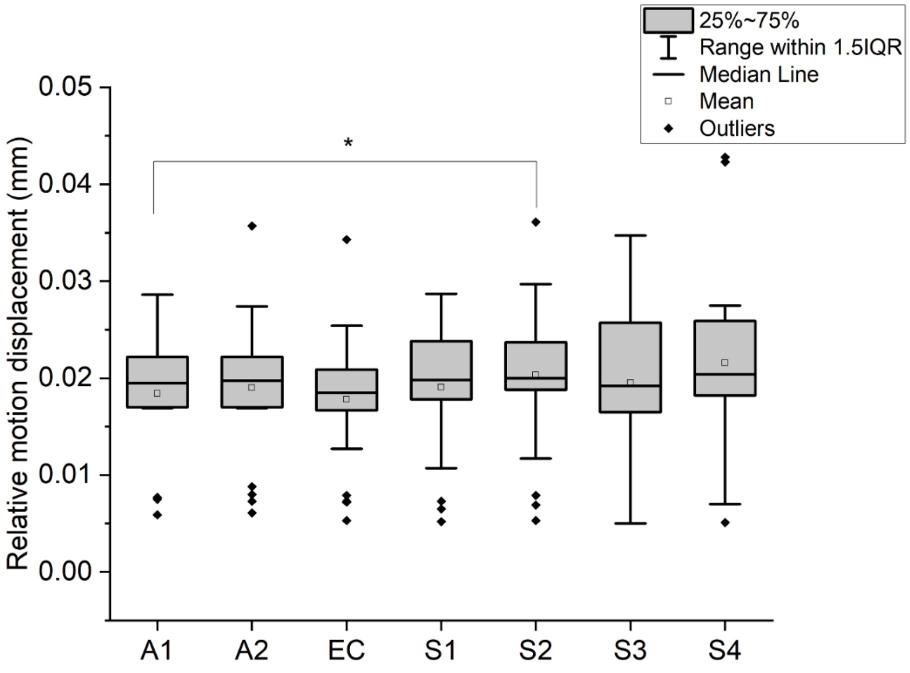


***Supplementary Figure 6. Relative mean head motion displacement (mm) in five-minute time segments in awake scans ($A_1$, $A_2$ and EC) and sleep scans ($S_1$, $S_2$, $S_3$ and $S_4$).*** *Relative head motion was slightly lower only in $A_1$ vs. $S_2$. * $p < 0.05$.*
